# Identifying *Clostridioides difficile*-inhibiting gut commensals using culturomics, phenotyping, and combinatorial community assembly

**DOI:** 10.1101/767715

**Authors:** Sudeep Ghimire, Chayan Roy, Supapit Wongkuna, Linto Antony, Gavin Fenske, Abhijit Maji, Mitchel Chan Keena, Andrew Foley, Joy Scaria

## Abstract

A major function of the gut microbiota is to provide colonization resistance, wherein pathogens are inhibited or suppressed below infectious level. However, the fraction of gut microbiota required for colonization resistance remains unclear. We used culturomics to isolate a gut microbiota culture collection comprising 1590 isolates belonging to 102 species. Estimated by metagenomic sequencing of fecal samples used for culture, this culture collection represents 50.73% of taxonomic diversity and 70% functional capacity. Using whole genome sequencing we characterized species representatives from this collection, and predicted their phenotypic traits, further characterizing isolates by defining nutrient utilization profile and short chain fatty acid (SCFA) production. When screened using a co-culture assay, 66 species in our culture collection inhibited *C. difficile*. Several phenotypes, particularly, growth rate, production of SCFAs, and the utilization of mannitol, sorbitol or succinate correlated with *C. difficile* inhibition. We used a combinatorial community assembly approach to formulate defined bacterial mixes inhibitory to *C. difficile*. When 256 combinations were tested, we found both species composition and blend size to be important in inhibition. Our results show that the interaction of bacteria with each other in a mix and with other members of gut commensals must be investigated for designing defined bacterial mixes for inhibiting *C. difficile in vivo*.

**IMPORTANCE:** Antibiotic treatment causes instability of gut microbiota and the loss of colonization resistance, allowing pathogens such as *C. difficile* to colonize, causing recurrent infection and mortality. Although fecal microbiome transplantation has shown to be an effective treatment for *C. difficile* infection (CDI), a more desirable approach would be the use of a defined mix of inhibitory gut bacteria. *C. difficile*-inhibiting species and bacterial combinations we identify herein improve our understanding of the ecological interactions controlling colonization resistance against *C. difficile*, and could aid the design of defined bacteriotherapy as a non-antibiotic alternative against CDI.

## INTRODUCTION

Normal functioning of the human gut requires a balanced interaction between our mucosal surface, diet, the microbiota, and its metabolic by-products. A major determinant of the gut homeostasis is the presence of a healthy, diverse commensal microbiota which prevents pathogenic bacteria from colonizing the gut or keeping their number below pathogenic levels. This function of the gut microbiome is called colonization resistance [1, 2]. Perturbations of the gut microbiome referred to as dysbiosis could result in the loss of colonization resistance[3]. Dysbiosis and loss of gut microbiome colonization resistance caused by antibiotics, for example, can predispose people to enteric infections. *Clostridioides difficile* infection (CDI) of the gut following antibiotic treatment is a clear demonstration of this phenomenon. *Clostridioides difficile* is a Gram-positive, spore forming anaerobe and is the leading cause of antibiotic induced diarrhea in hospitalized patients[4].

Antibiotic treatment of CDI often causes recurrence [5]. It has been shown that infusion of fecal microbiome from healthy people into CDI patients gut can resolve CDI and prevent recurrence [6, 7]. This procedure termed as fecal microbiome transplantation (FMT) has become a common treatment for CDI[8]. However, there has been concerns regarding the long term health consequence of FMT. Recently, there has been reports of weight gain [9] and mortality due to transfer of multi-drug resistant organisms following FMT[10].

The development of defined bacterial mixes from the healthy microbiota, which can resolve CDI, poses an alternative to FMT [11]. However, the exact number of species needed in an efficacious defined bacterial mix for CDI remains unknown, and was reportedly in the range of 10–33 species when defined bacteriotherapy was tested in a limited number of patients [12, 13]. A mix of spore-forming species tested in phase II clinical trials, despite its initial success, resulted later in recurrence [14]. The use of high-throughput anaerobic gut bacterial culturing, coupled with sequencing, improves the cultivability of the gut microbiota [15–17], facilitating the development of culture collections of gut commensals screenable for identification of species conferring colonization resistance, or an understanding of the ecological interactions stabilizing or destabilizing colonization resistance.

Here we report the cultivation, using culturomics, of 1590 gut commensals comprising 102 species from healthy human donors. We phenotyped and sequenced genomes of the representative species in this culture collection. We then screened the strains to identify species inhibiting *C. difficile*. A combinatorial community assembly approach in which 256 strain combinations were tested to identify the species interactions that improve or diminish *C. difficile* inhibition. Our results show that both species composition and interactions are both important determinants of *C. difficile* inhibition phenotype. Our approach and culture library—besides advancing the understanding of bacterial community interactions determining colonization resistance—could prove useful in other studies probing the role of gut microbiota in host health.

## RESULTS

### Single medium based culturomics retrieves high species diversity from donor fecal samples

In this study, we used metagenome sequencing to characterize the fecal microbiome composition of healthy human donors and culturomics to develop a strain library to identify *C. difficile* inhibiting species. As a first step, donor’s fecal samples were characterized using shotgun metagenome sequencing. To this end, fecal samples from six donors were sequenced individually and also after pooling in equal proportion. We used high sequencing depth for the metagenome sequencing. Collectively, the datasets from all samples constituted 48.9 Gb of data. To determine the taxonomic composition and diversity of the samples, Simpson dominance index (D), Shannon diversity index (H) and Shannon equitability index (E_H_) were calculated for individual samples as well the pooled material. This analysis revealed that all donor samples were similar in diversity indices and pooling the samples in equal proportion maintained the overall population structure of the individual samples (Figure 1A). The donors in this study were recent migrants to the United States from Asia and were expected to have a high proportion of Prevotella (Prevotella enterotype) in the gut microbiome. Consistent with this expectation, taxonomic diversity of the samples in phylum level (Figure 1B) showed the dominance of Bacteriodetes while the most abundant genus was Prevotella (Figure 1C).

**Figure 1:**
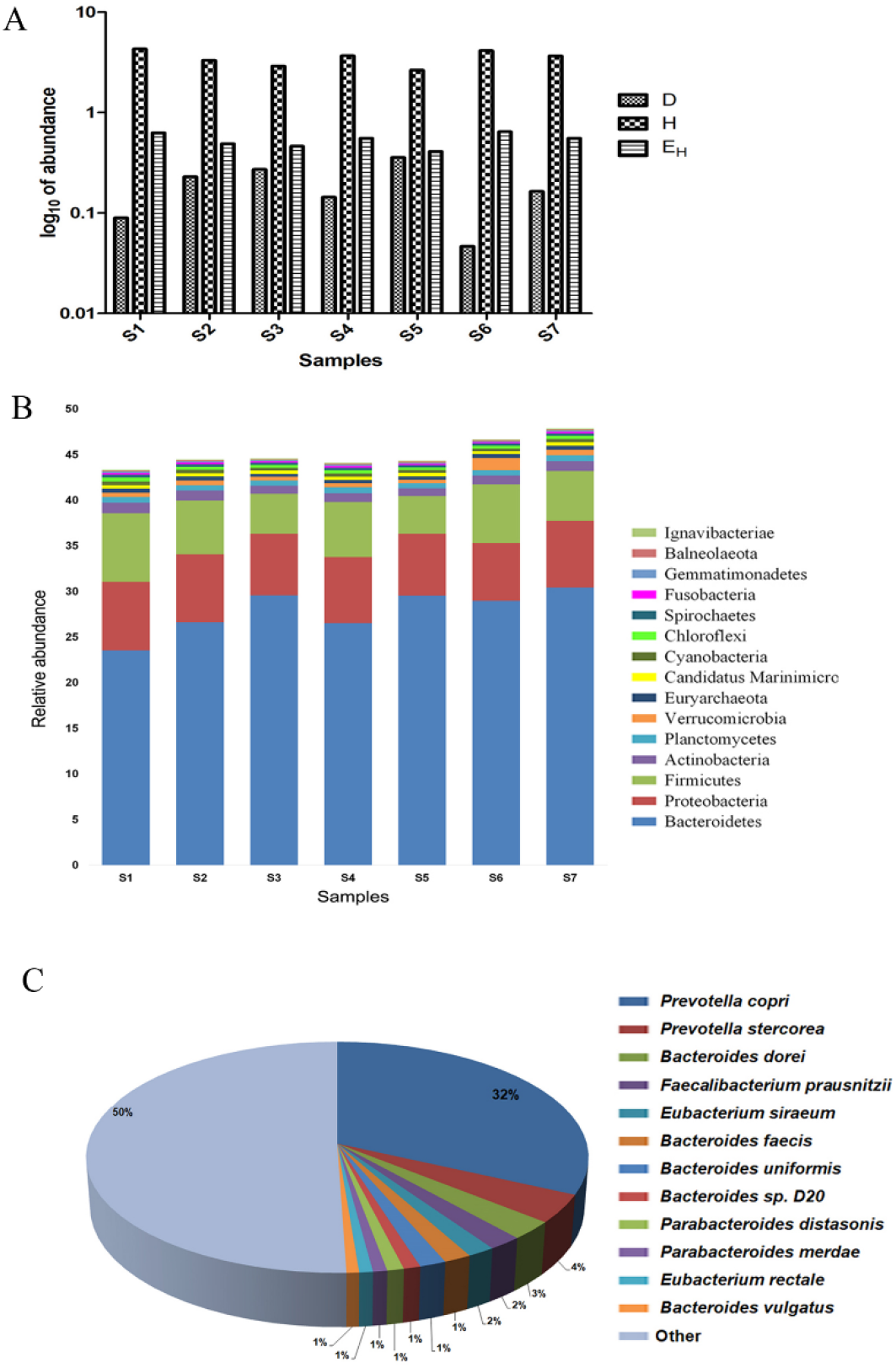
Gut microbiome composition of donor’s fecal samples. **A.** Variations in the Simpson Dominance (D), Shannon Diversity (H) and Shannon Equitability (EH) indices calculated based on the raw reads taxonomic affiliations for individual fecal samples (S1–S6) and pooled fecal sample (S7). **B.** Phylum level distribution of bacteria for individual fecal samples (S1–S6) and pooled fecal sample (S7). We calculated relative abundance values as compared to the total identifiable reads from any dataset. **C.** Species-level distribution of the pooled donor fecal sample (S7). We performed taxonomic classification using Kaiju. We referred all species with under 1% abundance to “other” category.

Using culturomics, we developed a strain library from the pooled fecal samples. Previous studies have prevalently used assorted media to isolate gut bacteria [15]. For mechanistic studies to understand the microbial community interaction, strains need to be pooled in a single nutrient medium. While the approach of using different media conditions is useful in retrieving high diversity, strains isolated in various media conditions may not be able to grow in a single media. This would prevent the use of a culture library in community assembly studied in a universal medium. To avoid this problem, we used a modified Brain Heart Infusion (mBHI) as the base medium for culturing. We isolated several strains from nine species from the mBHI medium. We reasoned that if the species that were growing fast in the mBHI medium is suppressed, additional species diversity could be isolated using the same medium. We therefore supplemented mBHI with different combinations of antibiotics selected to suppress the formerly dominant strains, also using heat shock and chloroform treatment to select for spore-forming species. We used 12 conditions for culturing (Supplementary Table S1), selecting 1590 colonies from these conditions. We determined strain species identity using MALDI-ToF, and 16S rRNA sequencing (Supplementary Table S2). We thus isolated 93 more species from the same sample, increasing the total diversity in our strain library to 102 species (Figure 2). In Figure 3A we present the frequency of each species isolated in each culture condition. We further examined whether our approach could isolate high- and low-abundance species from the sample. We therefore sequenced the genome of one representative isolate from each species in our library (Supplementary Table 3), mapping the metagenome reads (Supplementary Table 4) against the individual species genomes. We mapped 37,669,789 reads obtained from the pooled sample to species whole genomes (Figure 3B), matching 19,109,642 reads—50.73% of the total metagenomic diversity—to the metagenomic reads. Our single medium-based culturomics method was able to isolate about 50% of the pooled culture sample diversity. We mapped about 20%reads against the *Prevotella copri* SG-1727 genome isolated from six different conditions— unsurprising, as the fecal donors belonged to the Prevotella enterotype. However, we isolated low-abundance species such as *Olsonella umbonate*, for which only 0.5% reads were mapped, in eight different conditions (Figure 3B). We isolated both low- and high abundance species with our technique, and used metagenome binning to estimate the number of species missed by our method. Matching metagenome bins against cultured species genomes, we found 50 matching culture isolates and 33 bins matching none, indicating that our method failed to cultivate those metagenome bins (Supplementary Table 5).

**Figure 2:**
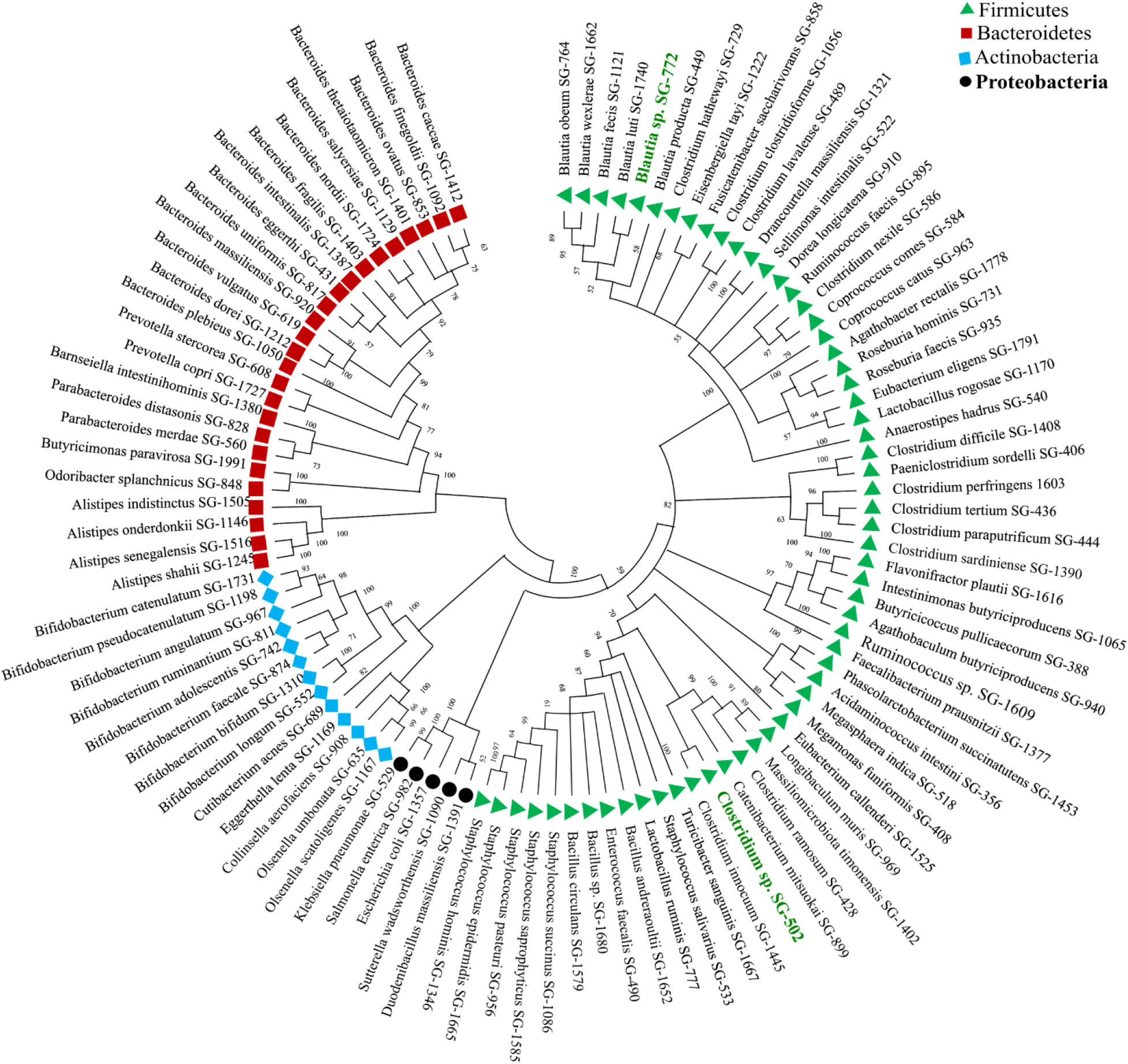
Neighbor-Joining tree of the full length 16S *rRNA* gene sequences of 102 cultured species isolated using single medium from the pooled donor fecal sample. We computed the evolutionary distances using the Jukes-Cantor method, presented in the units of the number of base substitutions per site in MEGA X. Symbols and colors represent four different bacterial phyla, as referred to the legend. We have highlighted putative novel species (n = 2) with “green” text.

**Figure 3:**
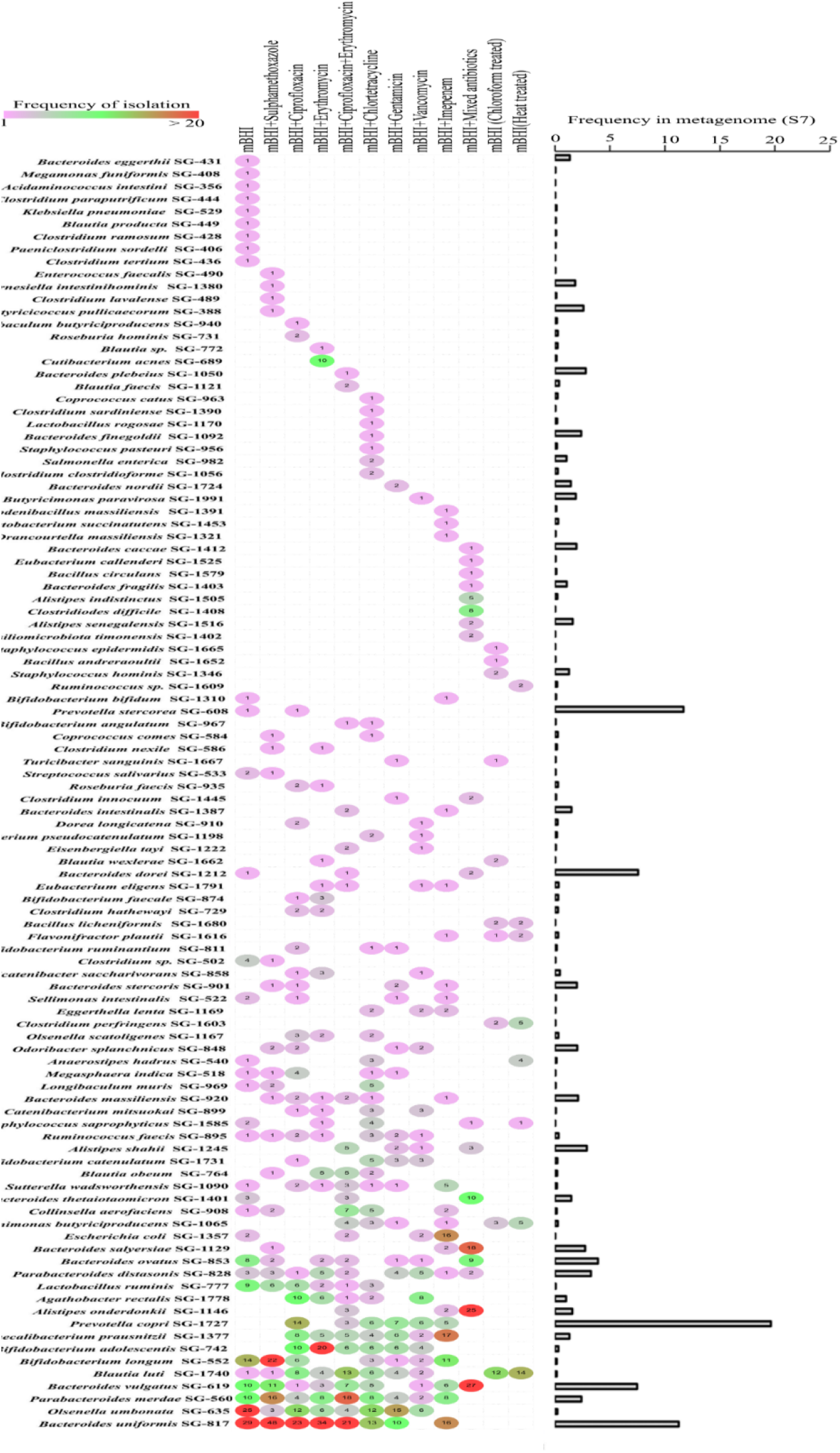
Efficiency of species retrieval in the culture conditions tested. **A.** Frequency of isolation of 102 species recovered. **B.** Prevalence of individual species as found from read mapping against donor fecal metagenome.

### Single medium based culturomics retrieves most frequent gut bacteria in human populations and increases gene repertoire in the integrated gene catalog of the gut microbiome

The availability of high-quality metagenome data from many human samples facilitates the identification of frequently present gut bacterial species in healthy populations. A recent study defined 71 bacteria present across 2144 human fecal metagenomes [18]. To quantify the number of these present in our culture library, we matched the whole genomes of 102 bacterial species and 33 metagenome bins against these 71 frequent species [18], finding our culture library contained 65 of these most frequent bacteria (Supplementary Table 5). To determine the relationship between the gene repertoire in our culture library and the integrated gene catalog (IGC) of the human microbiome, we compared the two datasets. For the comparison against IGC, a non-redundant gut microbiome gene set of 9.879 million genes generated using 1,267 human gut metagenomes from Europe, America and China [19], we generated a non-redundant gene set from our cultured genomes and the sample metagenome used for culturing. We created 984,515 open reading frames (ORFs) from sample metagenomes and 285,672 ORFs from cultured species genomes (Supplementary Figure 1A and Supplementary Table 6), and mapped them against the IGC. Of 285,672 ORFs obtained from cultured genomes, we matched 208,708 ORFs (73.05%) to the IGC, while 572,437 ORFs (58.14%) of 984,515 ORFs obtained from the sample metagenome matched the IGC (Supplementary Figure 1B and 1C). In other words, the IGC lacked 26.94% of the ORFs from the cultured isolates and 41.85% of the ORFs from the sample metagenome. This shows the potential for expansion of the IGC if more Prevotella enterotype donor fecal samples, such as those used in our study, are sequenced. Our results also show that sequencing cultured species genomes identifies genes otherwise missing in metagenome sequencing because of low depth or assembly issues.

### A large number of species in the culture library inhibits C. difficile in vitro

A healthy microbiota suppresses pathogen growth in the gut. To identify *C. difficile-inhibiting* species in our culture library, we screened it against *C. difficile* using a co-culture assay. Slow-growing strains would be outcompeted by *C. difficile*.We therefore used 82 moderately- or fast-growing species in the co-culture assay. When tested, 66 species inhibited *C. difficile* to varying degrees (Figure 4). In this screen, *Bifidobacterium adolescentis* strain SG-742 was the most efficient inhibitor. Furthermore, all *Bifidobacteria* inhibited *C. difficile*, signifying their importance in colonization resistance to this pathogen. The *Lachnospiraceae* family were major inhibitors. In a surprising result, 16 species in our co-culture assay increased the growth of *C. difficile* (Figure 4), a finding of clinical significance, in that a high abundance of these species may confer a higher risk of CDI.

**Figure 4:**
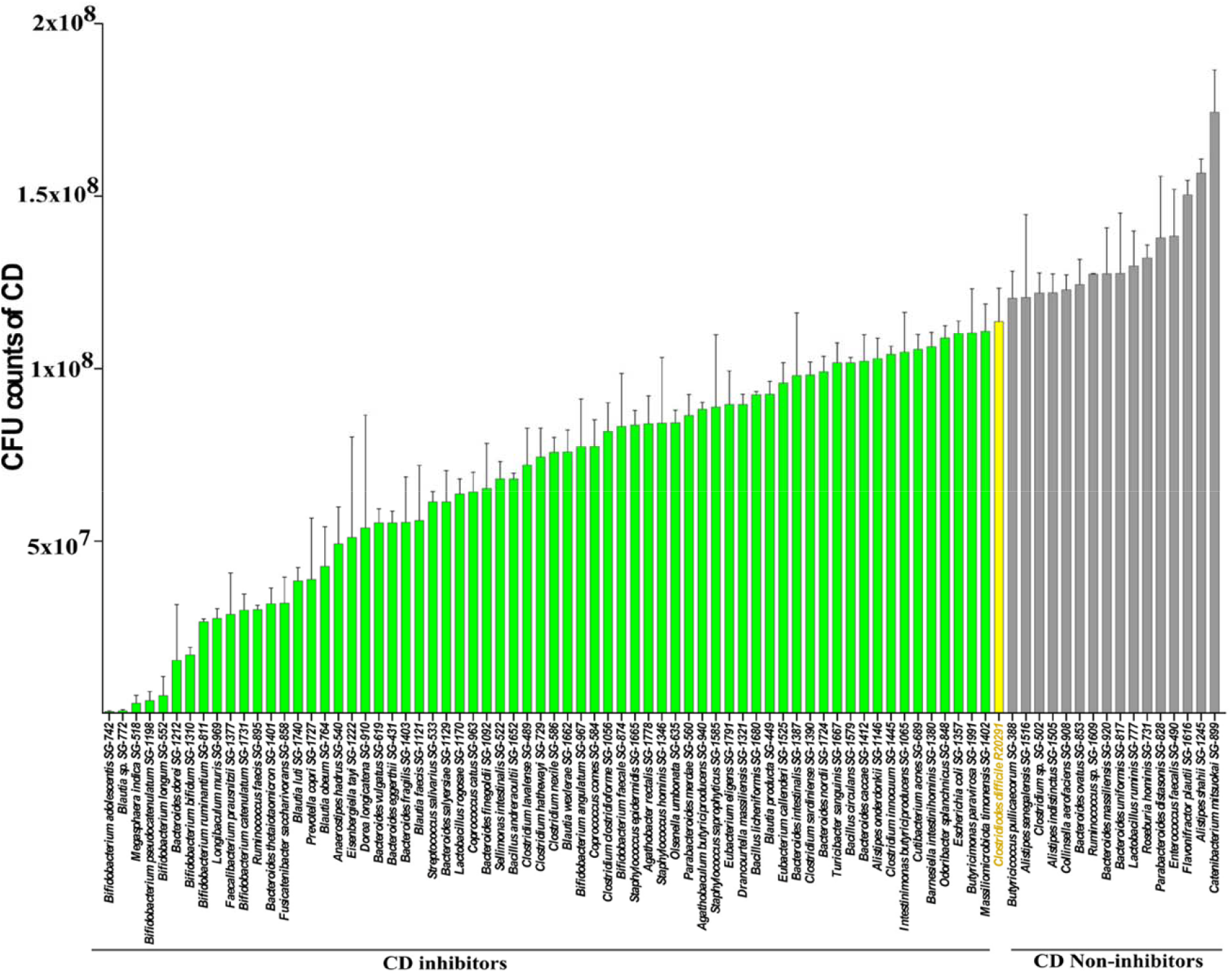
Inhibition of *C. difficile* by individual species *in vitro*. *C. difficile* CFU counts were plotted for every individual co-culture assay performed for 82 test species in triplicate. Error bars represent standard deviations of the three independent experiments for each inhibition assay for individual species. “Green,” “yellow” and “gray” bars represent CFU counts of *C. difficile* for inhibitors, control *C. difficile*, and non-inhibitors, respectively in co-culture.

### Most inhibitors are acetate or butyrate producers

Gut bacteria metabolize diverse substrates to produce short chain fatty acids (SCFAs) in the gut [20, 21]. SCFAs—particularly butyrate—act as gut epithelial immune modulators, energy sources for host intestinal cells, and pathogen-inhibitors [22, 23]. To determine the relationship between SCFAs produced and *C. difficile* inhibition, we estimated the SCFAs produced by all species used in the co-culture assay (Figure 5). Our results show that the strains produce mainly acetate (Figure 5A, Supplementary Table 7). Comparing all strains at phylum level using the non-parametric Kruskal–Wallis test, we found Actinobacteria and Bacteroidetes to yield the most acetate (Figure 5B). Firmicutes produced significantly higher butyrate than Bacteroidetes (Figure 5C). Other SCFA production did not differ significantly (Figure 5D & 5E). The majority of high acetate- or butyrate producers were *C. difficile*-inhibitors.

**Figure 5:**
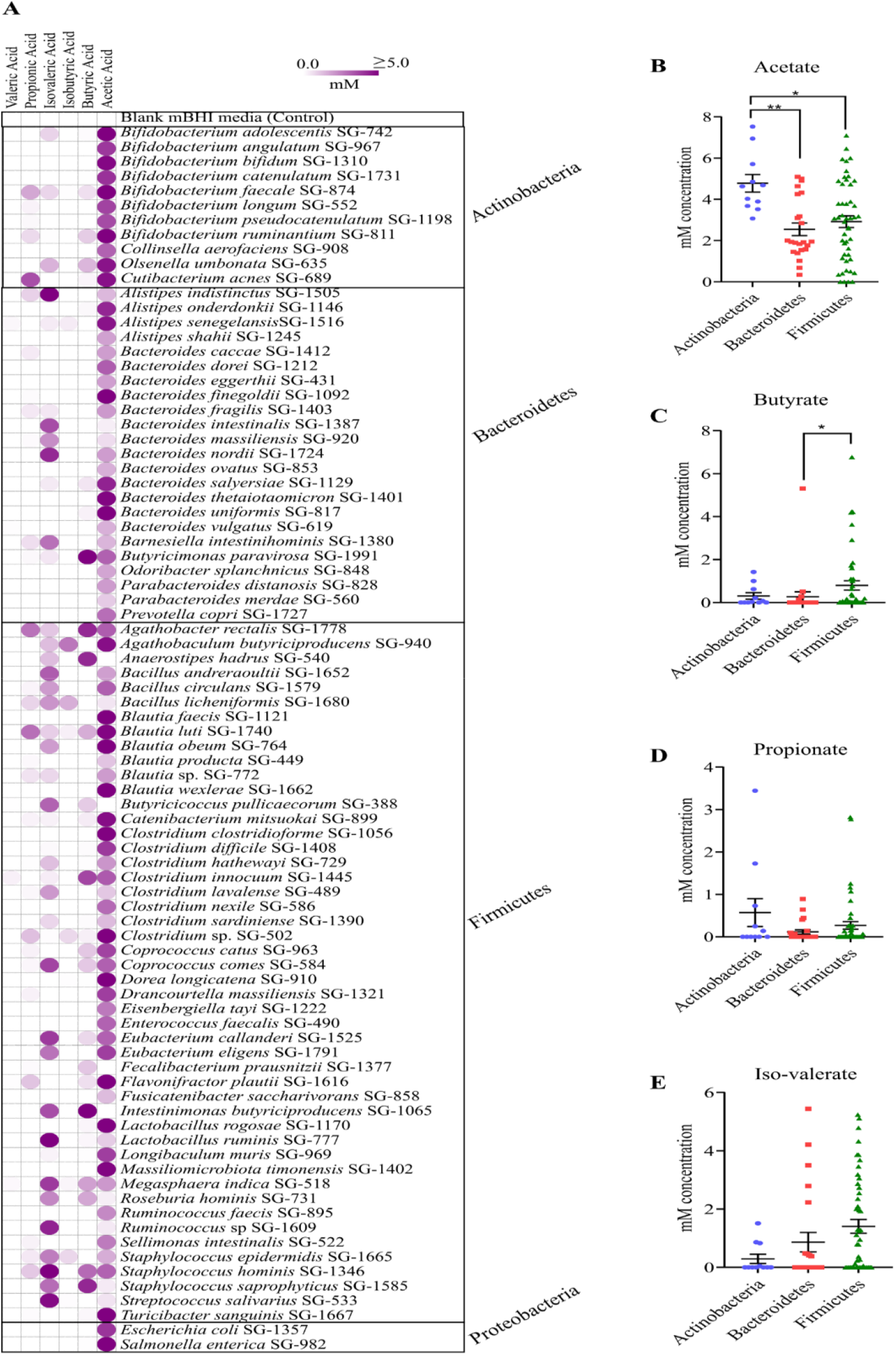
Short chain fatty acid (SCFA) produced by cultured species. **A.** SCFA production by 82 species used for inhibition assay against *C. difficile*. We measured acetate, propionate, isobutyrate, butyrate, isovalerate and valerate using gas chromatography, expressed in mM concentration. The figure represents mean SCFA measurements from duplicate samples. Color scale bar: “white”: no SCFAs production; “dark”: ≥5 mM SCFA production. **B.** Acetate, **C.** butyrate, **D.** propionate and **E.** isovalerate production in three major phyla from 80 bacteria when oriented according to phyla (Kruskal–Wallis tests, p<0.05). SCFAs levels for two members of Proteobacteria phyla are not shown in B, C, D, and E.

### Relationship between nutrient utilization, C. difficile inhibition and species prevalence in patientpouplations

Commensal species suppress pathogens in the gut chiefly by competing for nutrients [24–26]. The Biolog AN MicroPlate™ test panel provides a standardized method to identify the utilization of 95 nutrient sources by anaerobes. Investigating the relationship between nutrient utilization and pathogen inhibition, we determined the nutrient utilization of all species in our library. *C. difficile* is known to exploit mannitol, sorbitol or succinate to invade and produce infection in the human gut [27, 28]. Consistent with this observation, we found 27 *C. difficile*-inhibitors that utilized all three nutrients as carbon sources (Figure 6, Supplementary Table 8). We hypothesized that if the *C. difficile*-inhibitors we identified were active in *C. difficile* suppression *in vivo*, their abundance would be reduced in the gut during antibiotic treatment and CDI. Hence, we determined the abundance of the top 16 (top 25%) *C. difficile*-inhibitors in our library from a study of healthy and CDI patient gut microbiomes [29]. We determined the frequency of the top 16 *C. difficile*-inhibiting isolates in the following groups of human gut metagenomes; (a) CDI patients, (b) antibiotic-exposed but no CDI patients, and (C) no antibiotic exposure and no CDI (healthy) people. Consistent with our hypothesis, the top 16 *C. difficile*-inhibitors in our screen constituted about 20% of total abundance in the heathy microbiome, but were in low-abundance in both antibiotic-treated and CDI patient gut samples (Figure 7A). Interestingly, *Prevotella copri* (SG-1727)—among the most abundant species in the donor samples—was among the species depleted during antibiotic treatment and CDI (Figure 7B).

**Figure 6:**
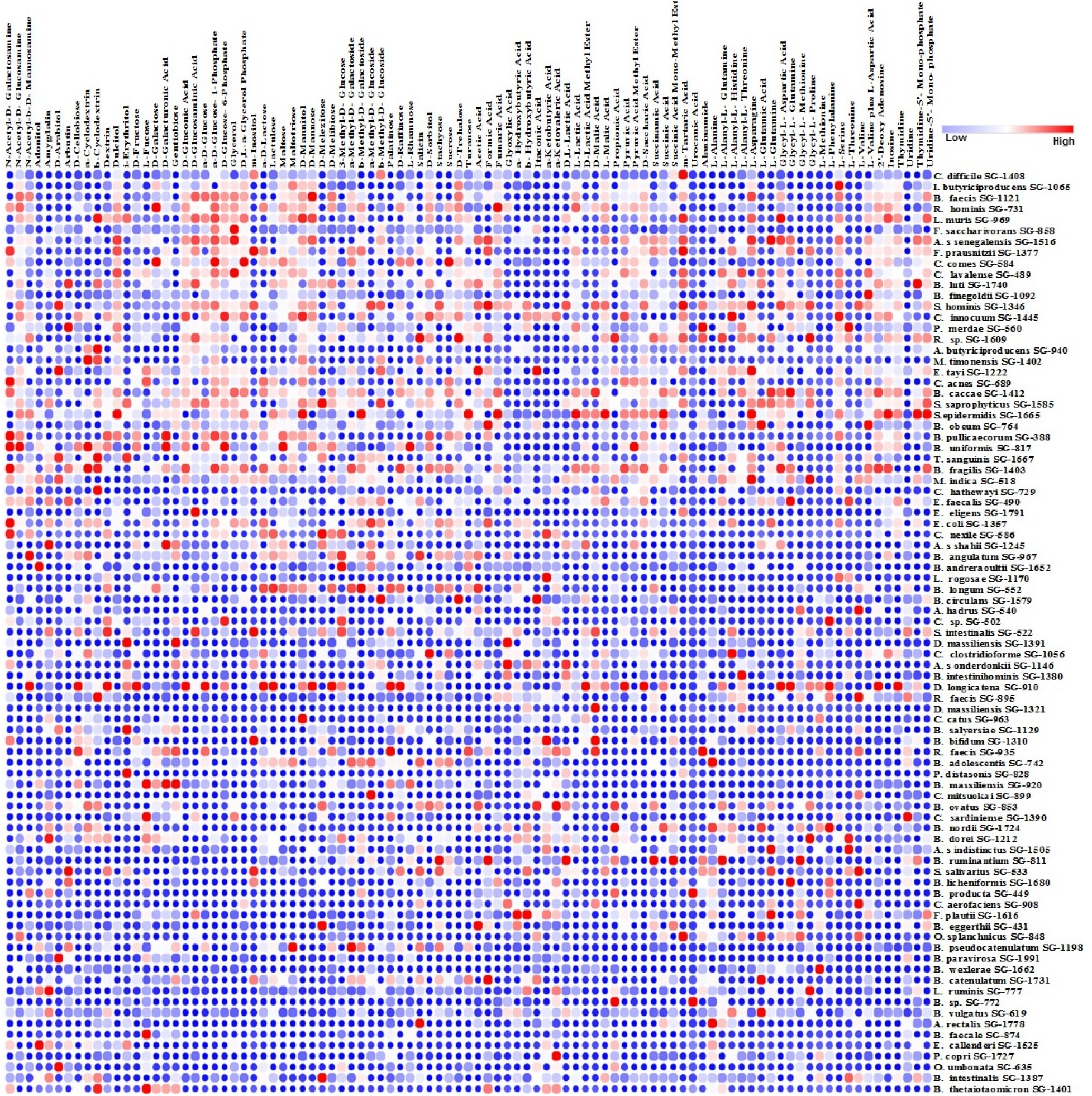
Comparison of nutrient utilization of 95 nutrient sources by the cultured species against *C. difficile*. The heatmap represents two independent substrate utilization tests normalized against control. Columns and rows represents nutrients and strains respectively. We considered growth of ≥ 20% in any substrate as compared to the control as positive. “Blue,” “white” and “red” represent high, medium and no utilization of the carbon source, respectively. Top row shows nutrient utilization of *C. difficile*. All other strains are arranged in descending order of nutrient utilization similarity when compared to C. difficile. We used nearest neighbor clustering based on the Pearson correlation to identify nutrient utilization similarity of other strains when compared to *C. difficile*.

### Analysis of genotype–phenotype relationships

To identify other phenotypes not covered in our phenotype assays and to link phenotype with genotype, we used Traitar [30] to predict 67 phenotypes from the genomes of all species in our library. The substrate utilization phenotypes based on the Traitar prediction mostly matched the Biolog phenotypes (Figure 8A). The first two clusters (green and red, lower Figure 8A) comprised mostly the pathogen-inhibitors tested here. The defining traits in these clusters related to sugar hydrolysis, mostly matching the Biolog phenotype data. The other two clusters (sky blue and mixed) comprised mostly pathogenic species and slow growers. Notable traits for pathogen clusters were catalase activity, beta hemolysis activity, growth in glycerol and high osmo-tolerance (Figure 8A). To further differentiate these traits, we performed principal component analysis (PCA) on the Traitar data. The PCA plot (Figure 8B) revealed four distinct clusters (*C. difficile*-inhibitors, -non-inhibitors, pathogens, and slow-growing strains) that explained 67.1% total variance in the first two principal components. Slow-growing strains clustered furthest from *C. difficile*-inhibitors—logically for any colonization-resistant bacteria, as a fast growth rate would more likely outcompete the pathogen.

To understand the genomic basis of colonization-resistant and pathogen-inhibiting strain genomes, we used KEGG modules—characteristic gene sets that can be linked to specific metabolic capacities or other phenotypic features of a genome. We identified 515 modules in the sample metagenome and 476 modules in the strain genomes. 432 modules were common between our sample metagenome and our strain genomes, demonstrating our ability to retrieve 77.28% of the metabolic functional capacity of the fecal microbiome using our culture method. 82 modules unrecovered in our isolated species comprised pathways related to environmental information processing, several components of cell signaling, DNA replication and repair pathways, lipid metabolism, RNA and protein processing, and the ubiquitin system (Supplementary Figure 2, Supplementary Table 9). We then tried to identify differences in KEGG modules associated with pathogen-inhibiting and -non-inhibiting strain genomes, finding 26 modules present in *C. difficile*-inhibitors were absent in non-C. *difficile-inhibitors*. Some important modules absent in non-inhibiting species genomes were M00698 (multidrug resistance efflux pump BpeEF-OprC), M00332 (Type III secretion system), M00438 (Nitrate/nitrite transport system), and M00551 (Benzoate degradation). Further work is necessary to understand how these functional modules relate to colonization resistance.

### Design of defined mix of *C. difficile*-inhibitors

While single strain vs pathogen co-culture assay is informative in identifying pathogen inhibiting strains, the inhibition patterns are likely to change when inhibiting species interact as a community. These communities may express emergent properties difficult to predict from the individual members [30]. After defining the isolate phenotypes, we used a combinatorial assembly of bacteria from our culture collection to design a tractable mix of *C. difficile*-inhibiting isolates. In the first set of experiments, we mixed 15 inhibiting species in equal proportion and tested them against *C. difficile* using the co-culture assay used for individual strains. Table 1 describes the overall properties of these isolates, selected based on the criteria of the medium pH not dropping below 5.6 after 24 h growth, and representation of overall taxonomic diversity at the family level. Investigating how changes in the mix composition could affect the inhibition capacity, we removed one or two species at a time from the mix of 15 species to create additional mixes, thus testing 121 mixes listed in Supplementary Table 10 against *C. difficile* in co-culture assay format. As shown in Figure 9, the removal of strains from the15-species mix had both positive and negative effects. When we removed species, several mixes were less effective than the 15-species mix, and the removal of two species increased the inhibition efficiency in a species dependent manner. Out of 121 mixes tested, mix number 22, comprising the species listed in Supplementary Table 10, most effectively inhibited *C. difficile* growth (by 79.41%). This clearly demonstrates the dependence of inhibition efficiency on the species composition of the defined mix used in the co-culture assay. Seeking the minimum number of species necessary for an effective *C. difficile*-inhibiting mix, we performed another set of experiments in which either one or two species at a time were removed from the parent mix of 12 species. In this round, we tested 79 bacterial mixes comprising species listed in Supplementary Table 10 in the co-culture assay. As shown in Figure 9, removal of species mostly reduced inhibition efficiency when compared with the parent blend. Again, the efficiency was dependent on the species composition of the mix. We performed a third set of experiments to determine mixes comprising under ten species that would impact *C. difficile* inhibition efficiency. Removal of species from the 10-species parent set diminished inhibition efficiency overall; however, some mixes increased the growth of *C. difficile*. Since all the strains we used individually inhibited *C. difficile*, the enhancement of *C. difficile* growth by these set mixes clearly demonstrates that individual strain phenotype can be overridden by species community interactions. In this case, a set of *C. difficile*-inhibitory species, when mixed in a particular combination, increased *C. difficile* growth. Overall, our results demonstrate that new phenotypes masking the individual strain phenotype could emerge when microbial consortia are formed, and this emergent property needs to be taken into account while designing defined *C.difficile*-inhibiting bacterial mixes.

**Table 1–.** Strains used in the combinatorial mix assembly. *C. difficile* inhibtion efficiency of strains are denoted as; high inhibition = ***, moderate inhibition = **, and low inhibition = *.

| Strain | Family | *C. difficile* inhibition | pH after growth in mBHI | Number of carbon sources utilized |
| --- | --- | --- | --- | --- |
| *Bifidobacterium bifidum* SG-1310 | *Bifidobacteriaceae* | *** | 6.333 | 30 |
| *Bacteroides eggerthi* SG-431 | *Bacteroidaceae* | ** | 6.100 | 35 |
| *Bacteroides finegoldii* SG-1092 | *Bacteroidaceae* | ** | 5.713 | 62 |
| *Bacteroides vulgatus* SG-619 | *Bacteroidaceae* | ** | 5.917 | 70 |
| *Parabacteroides merdae* SG-560 | *Tannerellaceae* | ** | 5.913 | 74 |
| *Prevotella copri* SG-1727 | *Prevotellaceae* | *** | 6.013 | 39 |
| *Lactobacillus rogosae* SG-1170 | *Lactobacillaceae* | ** | 6.003 | 22 |
| *Clostridium nexile* SG-586 | *Lachnospiraceae* | ** | 6.307 | 32 |
| *Eubacterium eligens* SG-1791 | *Eubacteriaceae* | ** | 6.237 | 26 |
| *Blautia wexlerae* SG-1662 | *Lachnospiraceae* | ** | 5.733 | 34 |
| *Sellimonas intestinalis* SG-522 | *Lachnospiraceae* | ** | 6.103 | 49 |
| *Drancourtella massiliensis* SG-1321 | *Ruminococcaceae* | ** | 5.770 | 5 |
| *Meghasphaera indica* SG-518 | *Veillonellaceae* | *** | 5.640 | 75 |
| *Bacillus licheniformis* SG-1680 | *Bacillaceae* | ** | 5.927 | 35 |
| *Clostridium sp.* SG502 | *Erysipelotrichaceae* | * | 5.880 | 30 |

## DISCUSSION

We developed a gut commensal culture collection from healthy human donors and identified *C. difficile*-colonization-resistant strains. As human gut microbiome composition varies across populations, donor selection for culture is an important consideration. Depending on the proportion of *Bacteriodes* and *Prevotella*, the human gut microbiome has been classified into enterotypes [31]. The Asian and African populations—two-thirds of the human population—fall into the Prevotella enterotype. [32]. Information about the colonization-conferring species in Prevotella-dominant gut microbiomes is limited; we therefore chose recent Asian immigrants in the US as the fecal donors in this study. For culture, we used a pooled sample from six donors before culturing, which could have positive and negative consequences. Pooling can save substantial time and resources. For instance, after pooling we analyzed 1590 colonies; were this to be done individually, the number of analyzed colonies would have been 9540. Pooling, however, may distort the microbiome composition of individual samples, creating artificial population assemblages [33]. For gut microbiome ecology studies, a gut commensal culture collection isolated from fecal donors of similar gut microbiome composition could be more useful than that from many different people. The publicly available gut microbiota culture collection from the human microbiome project is isolated from 265 people [34]. Other similar culture collections are also isolated from at least over 100 donors [15, 17]. Although such collections are useful as reference strains, since the donors may have different microbiome compositions, the strains isolated may not form stable ecologies if mixed together. In contrast, our collection, from a limited number of donors having similar microbiome composition, may form a stable ecology when mixed, and could be more useful for studies to understand underlying interactions determining colonization resistance and other traits.

Two general approaches have been used previously for developing gut microbiota culture collections: the use of culturing samples in several—often up to 64—different nutrient media conditions, to isolate diversity [15, 35, 36]; or the use of a single medium, needing less time and resource but retrieving fewer species [37]. In understanding the types of bacterial communities responsible for suppressing pathogen growth in the gut, the assemblage of simple to complex bacterial communities from cultured strain libraries and testing of such consortia against pathogens is commonly performed. Although the use of different nutrient media is highly efficient in isolating the maximum number of species from gut samples, strains so isolated may not grow in a common nutrient medium, diminishing the utility of the strains in community assembly studies. We therefore used a single medium-based approach using mBHI for culturing of the fecal bacteria. As shown in Figures 2 and 3, we isolated 102 species—representing about 50% species diversity determined by sequencing the donor fecal sample—by adjusting the mBHI medium (Supplementary Figure 1)—comparable to other single medium-based approaches [37–39]. Furthermore, results in Supplementary Table 4 and Supplementary Figure 2 shows that the 50% diversity isolated represents over 70% functional capacity of the donor gut microbiome. According to the insurance hypothesis of microbiota function, more than one species performing the same function is recruited in an ecosystem to allow for functional redundancy [40, 41], possibly explaining the recovery of 70% function from 50% species diversity in our library.

Many strains in our culture collection shown in Figures 4 inhibit *C. difficile* at varying levels. Several phenotypes—particularly, growth rate, production of SCFAs, and the utilization of mannitol, sorbitol or succinate—correlated with the *C. difficile*-inhibitor phenotype, consistent with previous reports that restoration of depleted SCFAs in the gut resolved CDI [42, 43] and competition between *C. difficile* and commensals for nutrients and increased availability of mannitol, sorbitol or succinate allowed *C. difficile* invasion of the gut [27, 28]. Top inhibiting species in our collection were also depleted in the CDI patient gut, indicating their role in providing colonization resistance against *C. difficile* (Figure 7). The formation of many different defined bacterial mixes using these inhibitory strains may improve the inhibition capacity of the individual strains. Overall, we tested 256 defined mixes using the combinatorial community assembly approach. The combinatorial community assembly method presented in Figure 9 shows two important parameters defining the efficacy of the defined mixes—the number and type of species in the mix. Reducing the number of species in the mix from 15 to 12, did not diminish the overall inhibition capacity, but reducing the number to ten species did so. Furthermore, many of those mixes increased, rather than inhibiting, *C. difficile* growth. Clearly, adding too many species in a mix does not improve inhibition. The threshold of peak efficacy is 12 species in the conditions we tested. Our results also underline how, when species are pooled in a sub-optimal ecology, undesirable traits could emerge; strains in combination could produce new phenotypes not observed individually. Previous work to identify a defined mix of *C. difficile*-inhibiting bacteria also identified varying mix numbers. For instance, more than 20 years ago, Tvede and Rask-Madsen showed that infusion of a 10-bacterial mix into a patient’s colon could resolve CDI [12]. Another study in a small patient population found treatment with a 33-bacterial mix could alleviate CDI. In a mouse model, *Clostridium scindens* was a more efficient *C. difficile*-inhibitor when mixed with a defined pool of other commensal bacteria [44]. Likewise, a mix of six phylogenetically diverse bacteria alleviated CDI in a mouse model [45]. The complexity of the gut microbiota and its variations across populations make the design of defined bacterial *C. difficile*-inhibiting mix not simply a matter of mixing large numbers of diverse species. Since our results show that the *C. difficile*-inhibiting phenotype changes substantially depending on the microbial interaction, design of a defined bacterial mix requires a deeper understanding of how inhibiting species interact with themselves and the members of the total commensal community.

**Figure 7:**
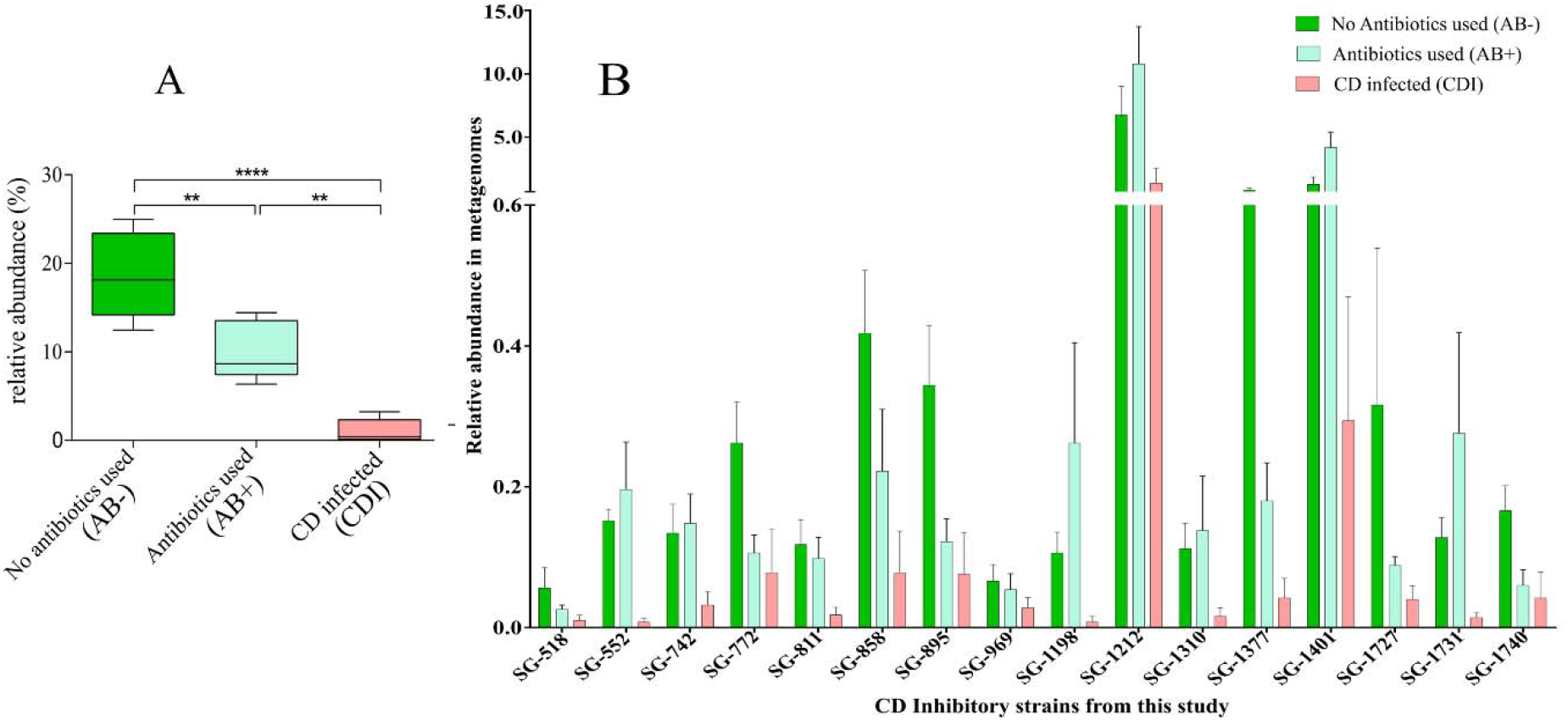
Frequency of the top 25% *C. difficile* inhibitors obtained in this study in the gut metagenome of *C. difficile* infected (CDI) and non-*CDI* patients. AB+ and AB- represent samples from non-*C. difficile* infected patients treated with antibiotics and non-*C. difficile* infected patients without antibiotic treatment, respectively. **A.** The combined abundance of top 25% CD-inhibitors (n = 16) from this study in CDI, AB+, and AB- metagenomes. **B.** Individual abundance of the same 16 strains in CDI, AB+, and AB- metagenomes. We obtained public metagenomes for CDI, AB+, and AB- from Milani et al., 2016.

**Figure 8:**
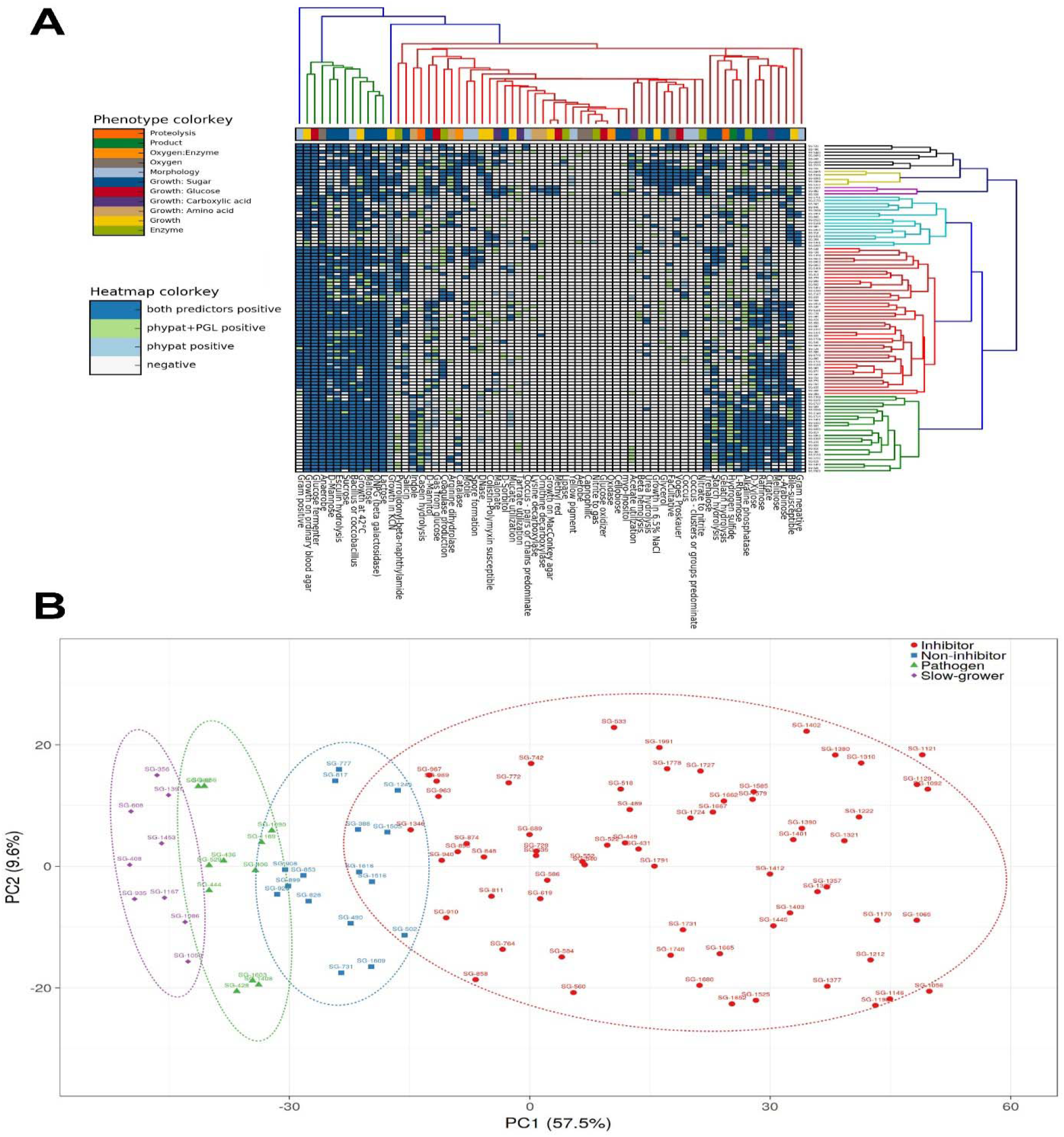
Prediction of phenotypes from the genomes of the 102 species in the culture library. **A.** Clustering of 102 species based on 67 traits predicted using the Traitar package. Each column represents one of 67 traits whereas rows represent 102 species from this study. The color scheme of the columns further depicts 11 phenotypic properties from proteolysis to enzyme production. The cluster colorings of the cladograms are tentative. **B.** PCA using the combined predicted traits from Pfam annotation. X- and Y-axes show Principal Component 1 & 2, explaining 57.5% and 9.6% of the total variance, respectively. Prediction ellipses are based on 0.95 confidences. Color scheme in the legends represents four different categories of isolates.

**Figure 9:**
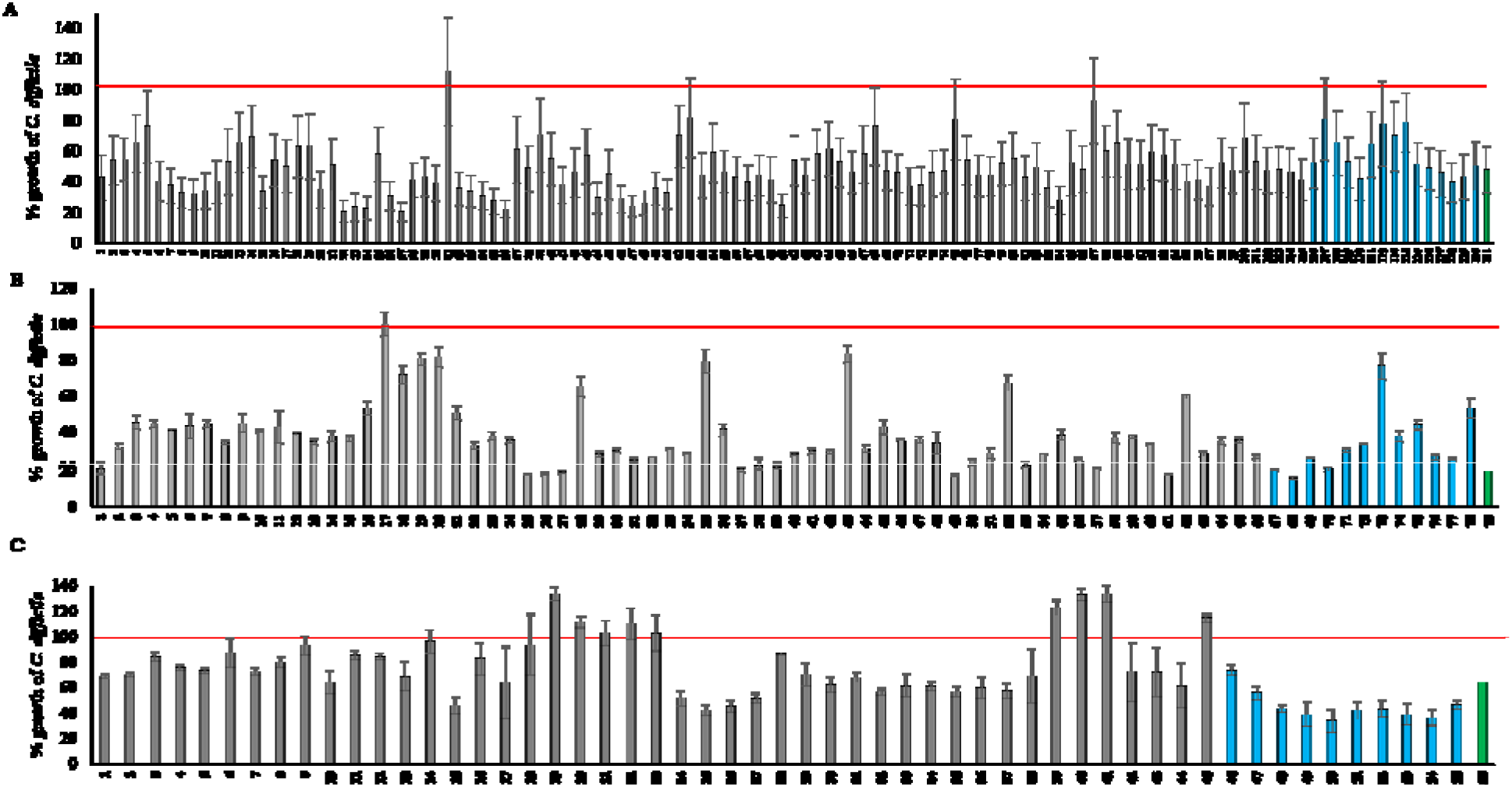
Inhibition of *C. difficile* by different consortia formed (Supplementary table 9) by combining individual *C. difficile*-inhibitors in different consortium sizes **A.** n =15, n =14, and n = 13; **B.** n = 12, n = 11, and n = 10; and **C.** n =10, n = 9, and n = 8. “Green,” “blue” and “gray” bars represent the parent blend, blend with one bacterial species removed at a time and blend with two bacterial species removed at a time, respectively. “Red” line represents *C. difficile* control growth. We normalized the growth of *C. difficile* in each consortium to control *C. difficile* growth, so as to obtain relative *C. difficile* growth (represented as % *C. difficile* growth) in each consortium condition *in vitro*. We performed each experiment with co-cultures, in triplicate, and error bars represent the standard error of the mean.

### Conclusion

Overall, we demonstrated that a high percentage of the cultivable fraction of gut microbiota from healthy human donors are *C. difficile*-inhibitors *in vitro*. Defined bacterial mixes can enhance the inhibition capacity of individual strains. However, depending on the ecology of the mix, new phenotypes could emerge. For instance, a mix of bacteria, rather than inhibiting, can increase the growth of *C. difficile*. In designing defined *C. difficile*-inhibiting bacterial mixes *in vivo*, the interaction of bacteria with each other in a mix and with other members of gut commensals demands investigation. The approach of combinatorial testing of strains with well-defined phenotypes used in the present study is a step in that direction.

## MATERIALS AND METHODS

### Fecal sample collection, culture conditions, isolate identification, and genome sequencing

We collected fecal samples with the approval of the Institutional Review Board (IRB), South Dakota State University. All procedures were performed following IRB guidelines. We collected fresh fecal samples from six healthy adult donors from Brookings, South Dakota, USA with no antibiotic consumption during the previous year. We transferred fecal samples to an anaerobic chamber within 10 minutes of voiding and diluted them 10-fold with phosphate-buffered saline, and mixed individual fecal samples in equal ratio to make the pooled sample for culturing. We used a modified brain heart infusion (mBHI) broth (ingredients listed in Supplementary Table 1) for culture. We used chloroform and heat treatment to isolate spore-forming species. We plated the 10^4^ dilution of the pooled sample on each of the media conditions described above in strictly anaerobic conditions. We selected 1590 isolates from all culture conditions and identified them using MALDI-ToF (MALDI Biotyper, Bruker Inc) against reference spectra (Supplementary Table 2). We identified isolates unidentified at this step using 16S rRNA gene sequencing, for which we prepared total genomic DNA using the OMEGA E.Z.N.A genomic DNA isolation kit (Omega Bio-tek, GA) according to the manufacturer’s protocol. We amplified full length bacterial 16s rDNA using the universal forward (27F) and reverse (1492R) primers, respectively, under standard PCR conditions. We sequenced the amplified DNA using the Sanger dideoxy method. We trimmed raw sequences generated for low-quality regions from either end and constructed consensus sequences from multiple primers using Genious [46]. Based on full length 16S rRNA gene sequence similarities, we determined the phylogenetic relationships of the isolates. For bacteria initially identified using MALDI-TOF, we extracted full length 16s RNA gene sequences from the genome using Barrnap (https://github.com/tseemann/barrnap) for phylogenetic tree creation. After obtaining the initial Neighbor-Joining tree, we performed heuristic searches for likelihood using the Nearest–Neighbor–Interchange and Close-Neighbor-Interchange branch swapping algorithms. Finally we created the Maximum Likelihood tree with a bootstrap of 1000 replicates using MEGA6 [47].

To further characterize the strain library, we selected representative isolates from each species for whole genome sequencing, extracting genomic DNA from overnight culture of the isolates using the OMEGA E.Z.N.A genomic DNA isolation kit (Omega Bio-tek, GA) according to the manufacturer’s protocol. We prepared sequencing libraries the Nextera XT kit, sequencing them using Illumina 2x 300 paired-end sequencing chemistry on the Miseq platform. We first filtered the raw reads for quality and sequencing adaptors with PRINSEQ [48], and then assembled them *de novo* using Unicycler [49], performing quality check of the assembly results with QUAST [50] and Bandage [51]. We performed gene calling using Prokka [52] with a minimum ORF length of 100 bp.

### Characterization of donor fecal samples using shotgun metagenome sequencing

We extracted total community DNA from 0.25 g of each donor fecal sample using MoBio Powersoil DNA isolation kit according to the manufacturer’s instructions. To enrich microbial DNA, using previously published protocol, we depleted host DNA in the isolated total DNA before sequencing [53]. We prepared the sequencing library from 0.3 ng of the enriched DNA using the Nextera XT library preparation kit (Illumina inc. San Diego, CA), performing sequencing following the protocols used for bacterial genomes described above. After quality correction, we removed human host reads using Bowtie2 v.1.1.2[54] and performed taxonomy assignment using Kaiju [55]. We searched reads against the proGenomes [56] reference database of protein sequences containing a non-redundant set from more than 25,000 genomes from every species cluster recovered by specI [57], using the NCBI non-redundant database for comparative analysis. Thereafter, we calculated the Simpson dominance index (D), Shannon Diversity Index (H) and Shannon Equitability Index (E_H_), using an assembly-based approach to characterize the donor fecal metagenomes. For this, we assembled reads *de novo* using metaSPAdes [58], specifically designed for assembly of complex metagenomic communities. For initial assembly, we error corrected reads using spades-hammer with default parameters, thereafter checking assembly results with MetaQUAST [59]. We removed contigs of less than 500 bp from the resultant datasets, and performed ORF predictions on the filtered contigs with MetaGeneMark [60] with a minimum length cutoff of 100 bp.

### Computation of gene repertoire and the functional analysis of isolate genomes and donor fecal metagenomes

To determine the gene repertoire in the isolate genomes and donor fecal metagenomes, we constructed a non-redundant gene catalog from our total data using cd-hit [61, 62] for comparison against the previously published integrated catalog of reference genes in the human gut microbiome [19]. After gene calling, we clustered the concatenated datasets from the culture library using cd-hit at >95% identity with 90% overlap. We checked these datasets using BLAT to avoid over-representation in the gene catalog. We then compared our gene dataset against the previously mentioned integrated catalog [19]. For functional mapping, we mapped amino acid sequences from all the individual datasets and clustered non-redundant sets against the EggNOG database v3.0 [63].

For gaining a better insight into the putative and hypothetical population genomes present within the donor fecal samples, but not isolated using the culturomics approach, population genome bin creation is considered superior to taxonomy assignment of the raw reads. We therefore constructed population genome bins from the metagenomes using MaxBin2), mapping back raw reads on the assembled contigs using Bowtie2 [64] for coverage information. We further analyzed all high-confidence bins with specI [57] for species cluster determination. To determine the abundance of isolated species from the pooled donor fecal samples, we measured the coverage by read mapping with Bowtie2 [64] at 95% identity level. For the functional analysis, we performed KEGG annotation for the ORFs obtained from the pooled donor fecal metagenome and the isolate genomes. We searched data from all comparisons for KO modules using GhostKOALA [65], and performed hierarchical clustering of the datasets to generate heat maps with R (http://www.R-project.org/). We used Traitar[66] under default parameters to predict 67 phenotypes from the whole genomes of all species in the culture collection.

### Phenotypic characterization of the isolated strains

To correlate the genomic features with phenotypes, we further characterized the strains for which genomes were sequenced by determining the following phenotypic properties:

#### Carbon source utilization

We determined the ability to utilize 95 carbon sources using Biolog Biolog AN MicroPlate™. Briefly, we grew strains on mBHI plates anaerobically for 24 h–48 h at 37^0^C. We used a sterile cotton swab to scrape cells from the plates and suspended them in AN inoculating fluid (OD_650_ < 0.02), using 100 μl of this suspension to inoculate AN MicroPlate™ in duplicate, and incubated them at 37^0^C anaerobically. We took OD_650_ readings at 0 h and 48 h post-inoculation and normalized the results for growth against water and 0-h OD_650_ values.

#### Production of SCFA

To analyze the SCFAs produced by isolates, we grew strains in mBHI for 24 hours in anaerobic conditions, and added 800 μl of the bacterial culture to 160 μl of freshly prepared 25% (w/v) m-phosphoric acid, and froze them at −80^0^C. We thawed samples and centrifuged them (>20,000×g) for 30 min. We used 600 μl supernatant for injection into the TRACE1310 GC system (ThermoScientific, USA) for SCFA analysis.

#### Identification of *C. difficile*-inhibiting strains

We used a co-culture assay in which pathogen and test strains were cultured together at a ratio of 1:9 to identify *C. difficile-inhibiting* strains in our library, using only those strains reaching OD600 of 1.5 after 24 h of growth in mBHI; 82 species met this criterion. We used *C. difficile* strain R20291 as the reference strain in the first assay. Briefly, we grew all test strains and *C. difficile* R20291 in mBHI medium anaerobically at 37^0^C and adjusted the OD_600_ to 0.5. The pathogen and the test were mixed together at a ratio of 9:1 and incubated for 18 hours anaerobically at 37^0^C. We then plated 10^-5^ & 10^-6^ dilutions onto *C. difficile* selective agar, using mono-culture of *C. difficile* R20291 as a positive control. We compared colony forming units (CFUs) enumerated from co-culture plates against the C*. difficile* R20291 control. In identifying *C. difficile*-inhibitors in healthy and CDI patients, we calculated the frequency of 16 top *C. difficile*-inhibitors in our collection by metagenome read mapping from a previously published dataset [29].

## Supplementary Figures

**Supplementary Figure 1:**
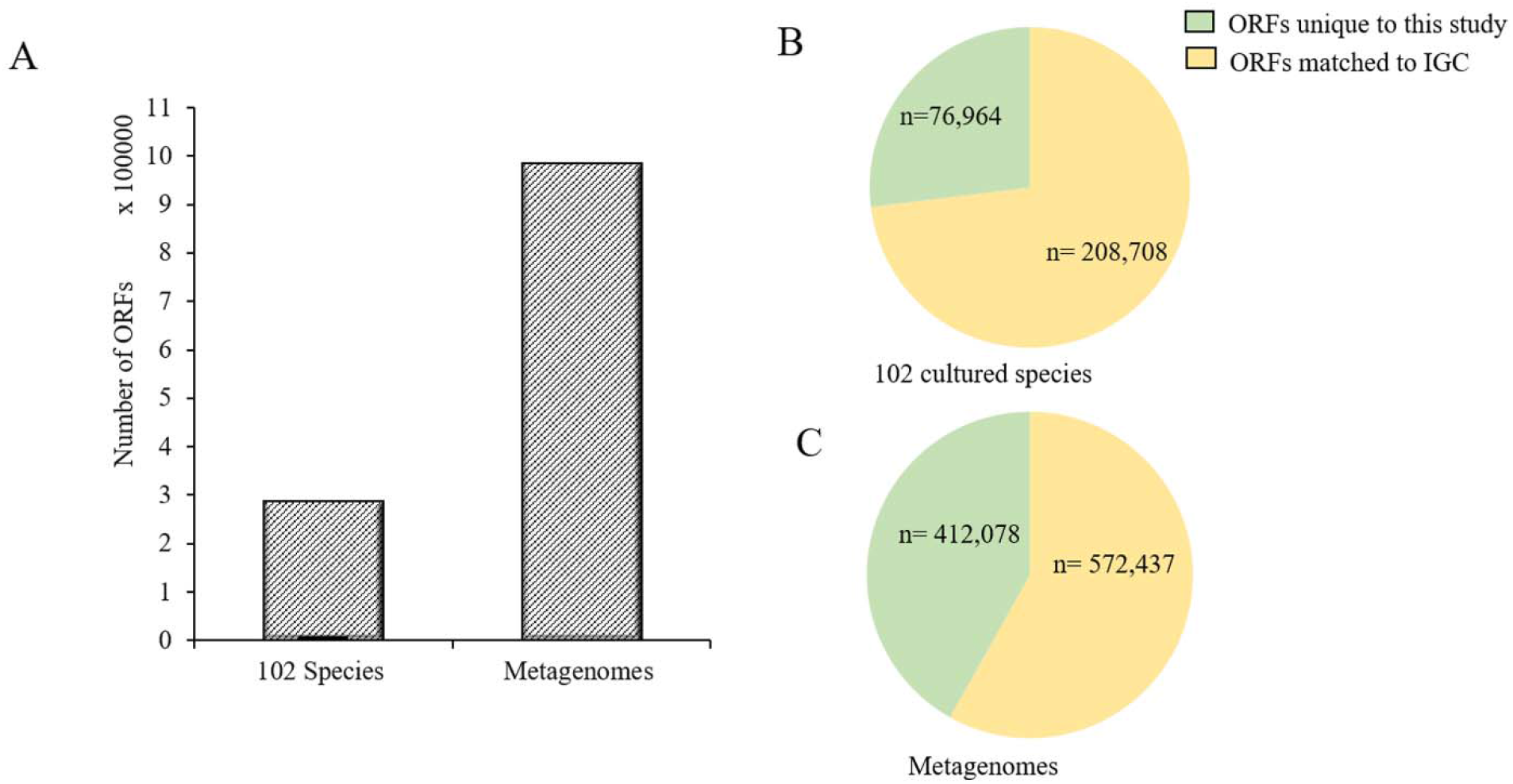
**A.** Numbers of non-redundant ORFs predicted in 102 cultured species and metagenomes (S1–S7). We generated the number of non-redundant ORFs at 95% identity cutoff for donor metagenomes and 102 isolates. **B.** Comparison of the non-redundant ORFs generated form 102 cultured species with the existing integrated human gene catalog (IGC). **C.** Comparison of the non-redundant ORFs generated from donor metagenomes with the existing IGC.

**Supplemtary Figure 2:**
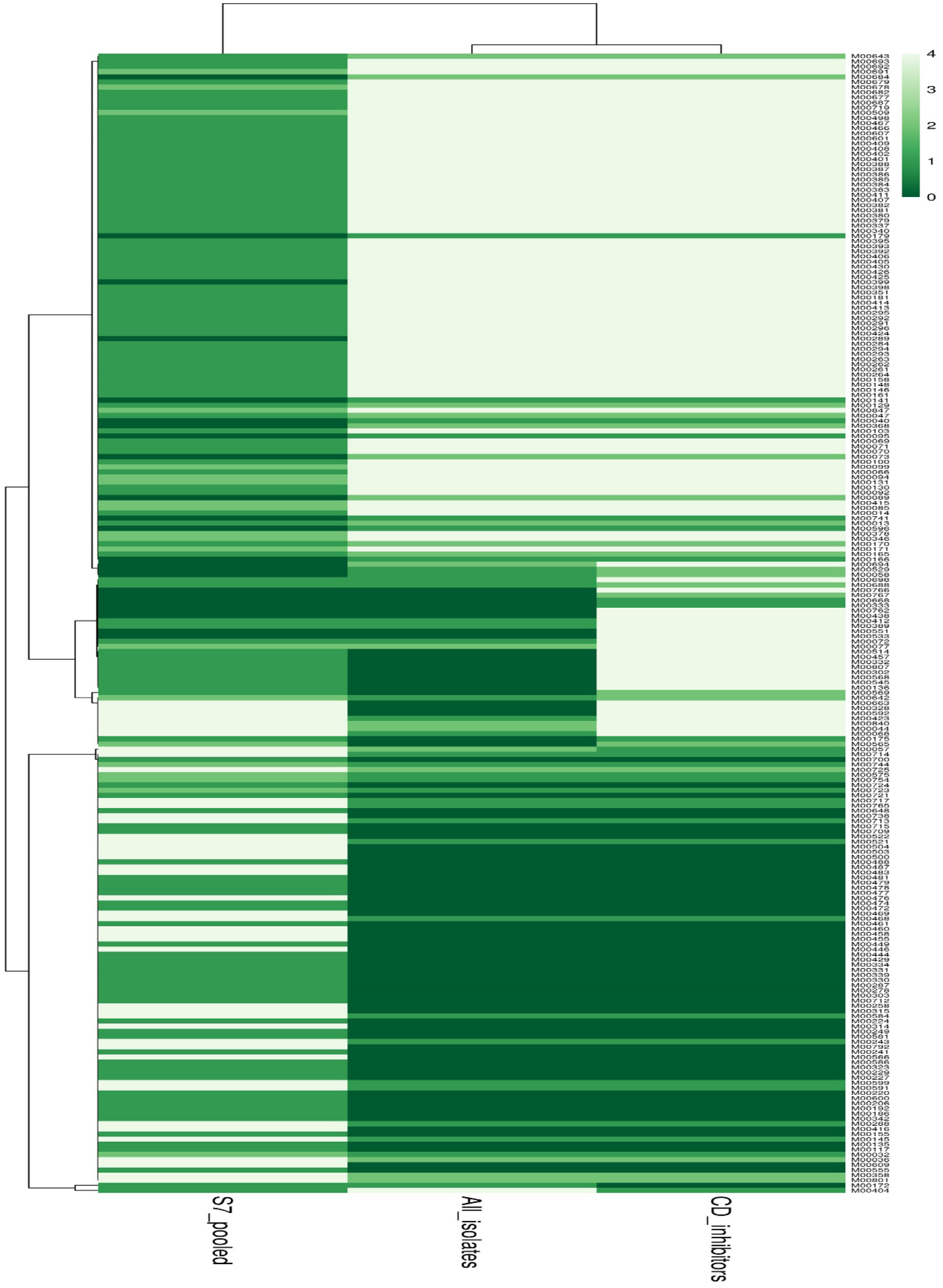
Hierarchical clustering of KEGG modules from fecal sample metagenome, all the 102 species (All_isolates), and subsets that were found to inhibit *C. difficile* R20291 (CD_inhibitors). We annotated predicted ORFs from the pooled donor (S7) metagenome and consortium of cultures for KO modules by searching against the KEGG database using Ghostkoala. We indicate the completeness of the KEGG modules by the color gradient where we refer to a complete module by “0” and the absence of a whole module by “4.” For detailed naming of the KEGG modules see Supplementary Table S6.

## Supplementary Tables

**Supplementary Table S1:** Composition of modified brain heart infusion (mBHI) medium along with antibiotic dosages used for isolation of individual isolates from pooled donor sample (S7).

**Supplementary Table S2:** Summary of identification strategy and taxonomic classification of all 1590 bacterial isolates used for the culturomics study.

**Supplementary Table S3:** Summary of metagenomic investigation of the six donors (S1–S6) and pooled donor (S7) fecal sample. We deposited all samples under the BioProject PRJNA494584.

**Supplementary Table S4:** Summary of metagenomic assembly of the six donors (S1-S6) and pooled donor (S7) fecal metagenomic samples used for this study.

**Supplementary Table S5:** Summary of the genomic features and sequence accession numbers for 102 bacterial species isolated using culturomics from the pooled donor fecal sample (S7). We deposited all samples under the BioProject PRJNA494608.

**Supplementary Table S6:** Comparison of the 102 species isolated in culture and high-quality bins obtained from metagenomes (S1–S7) from this study, with the most abundant 71 species and 20 species reported by Costea et al., 2017 and Froster et al., 2019, respectively. The numbers “1” and “0” denote presence and absence of the bacteria in the studies.

**Supplementary Table S7:** Differential presence of KEGG modules in pooled donor fecal sample (S7), all cultured 102 species and *C. difficile* R20291 (n = 66) inhibitors. The table provides supplementary data for Supplementary Figure 2, where 0 refers to a complete module and 4 refers to a module completely absent. 1 and 2 refer to number of absent modules.

**Supplementary Table S8:** Average short chain fatty acids (SCFAs) in mBHI medium for 82 species for which we performed an inhibition assay against *C. difficile* R20291.

**Supplementary Table S9:** Bacterial combinations formed to generate different consortium sizes A) n = 15, n = 14, and n = 13; B) n = 12, n = 11, and n = 10; and C) n = 10, n = 9, and n = 8. We formed consortia using the parent blend of 15 strains, 12 strains, and 10 strains, for three different experiments. For each experiment, we formed a parent blend, a blend with one bacterial strain removed at a time, and a blend with two bacterial strains removed at a time. “Green,” “blue” and “gray” colors represent the parent blend, the blend with one bacterial strain removed at a time, and the blend with two bacterial strains removed at a time, respectively. The blend numbers correspond to those in Figure 9.

